# Rapamycin alters the feeding preference for amino acids and sugar in female *Drosophila*

**DOI:** 10.1101/2024.09.10.611925

**Authors:** Guixiang Yu, Qihao Yang, Qi Wu

## Abstract

Pharmacological interventions targeting the aging process hold significant promise for improving the quality of life in the elderly and reducing healthcare costs. Rapamycin, in particular, has exhibited significant anti-aging and lifespan-extending effects across multiple model organisms. However, chronic rapamycin administration may also lead to various adverse reactions since it reshapes energy metabolism. Here, using *Drosophila melanogaster* as a model, we show that life-prolonging doses of rapamycin significantly modify animal feeding behavior. Long-term administration of rapamycin decreased protein preference in females while enhancing their sugar intake. Utilizing a chemically defined diet, we identified that changes in amino acid and sugar feeding preferences emerged as early as the second day of rapamycin treatment, preceding any detectable decline in fecundity. However, rapamycin-induced changes in macronutrient feeding preferences were not observed in males and sterile mutant females. Overall, our study suggests that the modification of feeding behavior could be a non-negligible side effect of rapamycin treatment, which is influenced by both sex and reproductive status.

## 1. Introduction

In light of the global demographic shift towards an aging population, pharmacological interventions targeting the biological mechanisms of aging are poised to significantly enhance the quality of life for the elderly and reduce healthcare costs. Over the past two decades, some drugs with anti-aging and life-prolonging activities have been identified in model organism studies, including rapamycin, metformin, spermidine and senolytic drugs ^1-4^. Among them, rapamycin has been demonstrated to increase lifespan in yeast, nematodes, fruit flies and mammals, primarily by modulating the conserved mTOR signaling pathway. The life-extending effect of rapamycin has been linked with marked improvements in multiple age-associated pathological phenotypes, such as cognitive and motor decline, as well as compromised intestinal barrier function^1,5-8^. In addition, studies in *Drosophila* and mice have shown that even short-term rapamycin administration is sufficient to delay aging and prolong life^9,10^.

Despite its powerful anti-aging effects, there are potential side effects associated with rapamycin use as an anti-aging drug. In murine studies, rapamycin treatment has been associated with metabolic disruptions, notably in glucose and lipid metabolism, leading to insulin resistance, hyperglycemia, and hyperlipidemia^11,12^. Its immunosuppressive properties further elevate the risk of infections^13^. Additionally, rapamycin has been shown to impair ovarian function in females and increase the incidence of testicular degeneration in males, thereby reducing reproductive capacity^1,6^. In young animals, rapamycin may also hinder normal growth and development^14^. Nonetheless, the potential behavioral side effects of rapamycin as an anti-aging agent remain inadequately understood. Given the prominent effects of rapamycin on animal reproduction and energy metabolism, which are important determinants of nutrient requirements, we investigated on the effect of rapamycin on animal feeding preferences in this study.

## 2. Methods

### 2.1 Fly stocks and husbandry

The wild-type stock Dahomey was collected in 1970 in Dahomey (now Benin). BL1,309(ovo[D1] v[24]/C(1)DX, y[1] w[1] f[1]) were obtained from the Bloomington Stock Center and backcrossed into wildtype *Dahomey* for eight generations. *ovo*^*D1*^ males were crossed with *Dahomey* females to obtain sterile *ovo*^*D1*^ infertile females. All stocks were maintained at 25°C on a 12-hr: 12-hr light:dark cycle at 60% humidity using 1SY food (7 g/L agar; 50 g/L sucrose; 100 g/L yeast. For all experiments, flies were reared at standard larval density in 1SY food and eclosed adults were collected over a 12-hr period ^15^. Flies were mated for 48 hr on 1SY food in all experiments before sorting into single sexes. 48 hr-mated flies were collected and transferred to 1SY or FLYaa holidic medium ^16^ for lifespan and fecundity experiment and maintained at 25°C on a 12-hr:12-hr light : dark cycle at 60% humidity.

### 2.2 Lifespan analysis

Flies were randomly allocated to the experimental food treatments and housed in plastic vials containing food at a density of 10 flies per vial, with 10 vials per condition (n = 100). Flies were transferred to a fresh food source every 2–3 days, during which any deaths and censors were recorded. Lifespan differences were assessed using the log-rank test.

### 2.3 Fecundity

The number of eggs laid in 24-hr periods was counted in all experiment. For each condition and each time point, 10 vials were counted. Each vial contained 10 flies. Egg-laying differences were assessed by unpaired t-test.

### 2.4 Binary choice EX-Q feeding assay

Feeding preference was measured using a modified apparatus based on the excreta quantification (EX-Q) feeding assay ^17^. Briefly, two foods were installed in each EX-Q tube, and 0.2% Erioglaucine disodium salt (Sigma Aldrich, 861146) was added to one of the foods. Flies were transferred to EX-Q tubes and kept for 24 hr at 25°C on a 12-hr:12-hr light : dark cycle at 60% humidity (10 flies/tubes, 20 tubes/treatment). After 24hr, flies were transferred out of the EX-Q tubes and the dye in excreta were then dissolved in 2 mL 0.1% PBST. The absorbance of the liquid sample was then measured at 630 nm (Molecular Devices, FlexStation 3) and used for food intake calculation according to a standard curve prepared from stock solutions of pure dye.

## 3. Results

### 3.1 Rapamycin decreases the feeding preference for yeast extracts in female *Drosophila*

Previous research has demonstrated that rapamycin significantly extends the lifespan of female fruit flies, with minimal impact on male lifespan^1^. Therefore, to investigate whether life-extending doses of rapamycin alter feeding behavior, we initially sought to verify if the optimal life-prolonging dose identified in prior studies would similarly extend the lifespan of female flies under our laboratory conditions. Consistent with previous studies, 50 mM of rapamycin in sucrose-yeast medium (comprising 10% yeast + 5% sucrose) significantly extended the lifespan of wild-type females and reduced their fecundity **(Figure 1A, B)**. Given that the nutritional requirements of egg-laying influence females’ protein consumption^18,19^, we further assessed the effect of long-term rapamycin exposure on females’ yeast extract and sugar preferences at various ages using the EX-Q feeding assay^17^ **(Figure 1C, D)**. At all four tested time points, the rapamycin-treated group displayed decreased yeast extract consumption and increased sugar intake compared to the control group, suggesting that rapamycin diminishes females’ preference for yeast extract **(Figure 1E, F)**. Intriguingly, even among 40-day-old females with notably low fecundity, rapamycin still considerably reduced the preference for yeast extract, implying that the drug’s impact on feeding preferences may not be solely attributable to reproductive inhibition.

**Figure 1.**
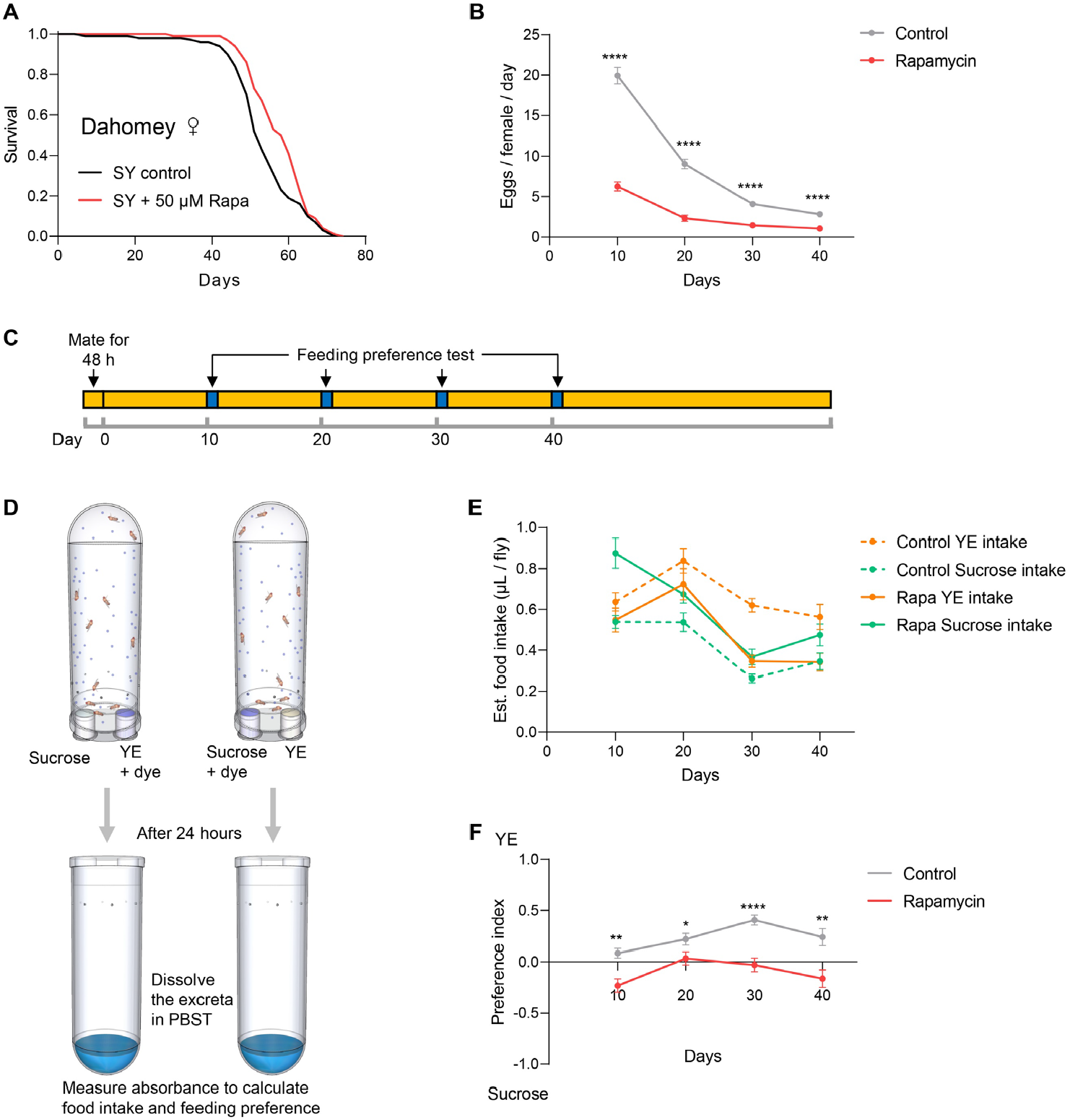
The life-extending dose of rapamycin changed the feeding preference of female flies for yeast extract and sucrose. (A-B) 50 mM rapamycin significantly increased the lifespan (A) and decreased fecundity (B) of Dahomey females. (C) Experimental timeline for rearing and feeding preference test in Dahomey females. The yellow bars represent the time period during which flies were fed with sucrose-yeast (SY) control medium or SY medium containing rapamycin, and the blue bars represent the time period at which feeding preference was tested. (D) Schematic diagram of the binary choice EX-Q feeding preference test. In two groups, 5% yeast extract medium and 5% sucrose medium were labeled with dye, respectively. The experimental flies were fed on binary choice medium for 24 hours, then the flies were transferred out, and the dye-containing excrement in the EX-Q tube was dissolved with PBST. Food intake and feeding preference were calculated by measuring absorbance. (E) Rapamycin decreased the food intake of yeast extract and increased the food intake of sucrose in females. (F) Rapamycin increased the female’s feeding preference for yeast extract. (For panel B and F, *p < 0.05,**p < 0.01, ****p < 0.0001, t-test)

### 3.2 Short-term rapamycin treatment alters amino acid and sugar feeding preference in a sex - and reproductively dependent manner

Since protein is the main nutrient component in yeast extract^20^, the feeding preference for yeast extract in many studies has often been interpreted as a preference for protein in *Drosophila*. Nonetheless, yeast extract also comprises a suite of other nutrients, including B vitamins, minerals, and nucleotides, some of which may influence *Drosophila* feeding behavior^20-23^. Therefore, to elucidate whether and how rapamycin impacts protein and sugar feeding preferences in *Drosophila*, we investigated the effects of short-term rapamycin treatment on the feeding preference for amino acids and sucrose using the FLYaa chemically defined medium^16^. We confirmed that a concentration of 5 mM rapamycin in the FLYaa medium significantly extended the lifespan of wild-type females and reduced their fecundity by the third day of treatment **(Figure 2A, B)**. To assess flies’ feeding preferences for amino acids and sucrose without inducing malnutrition from the absence of other nutrients, we provided flies with two food options: FLYaa medium devoid of amino acids and FLYaa medium devoid of sucrose, both of which contained all other essential nutrients, including cholesterol, vitamins, minerals, and nucleotides **(Figure 2C)**. We found that, beginning on the second day of rapamycin treatment, wild-type females showed a significant decrease in amino acid preference, characterized by reduced amino acid intake and increased sugar consumption **(Figure 2D, E)**. Importantly, this alteration in feeding preference occurred prior to the observed decrease in fecundity, which became significant on the third day of rapamycin exposure **(Figure 2B)**. In contrast, males and *ovo*^*D1*^ sterile mutant females did not exhibit any changes in feeding preferences for amino acids or sucrose **(Figure 2F-I)**, suggesting that rapamycin’s effects on feeding behavior are contingent on both sex and reproductive status.

**Figure 2.**
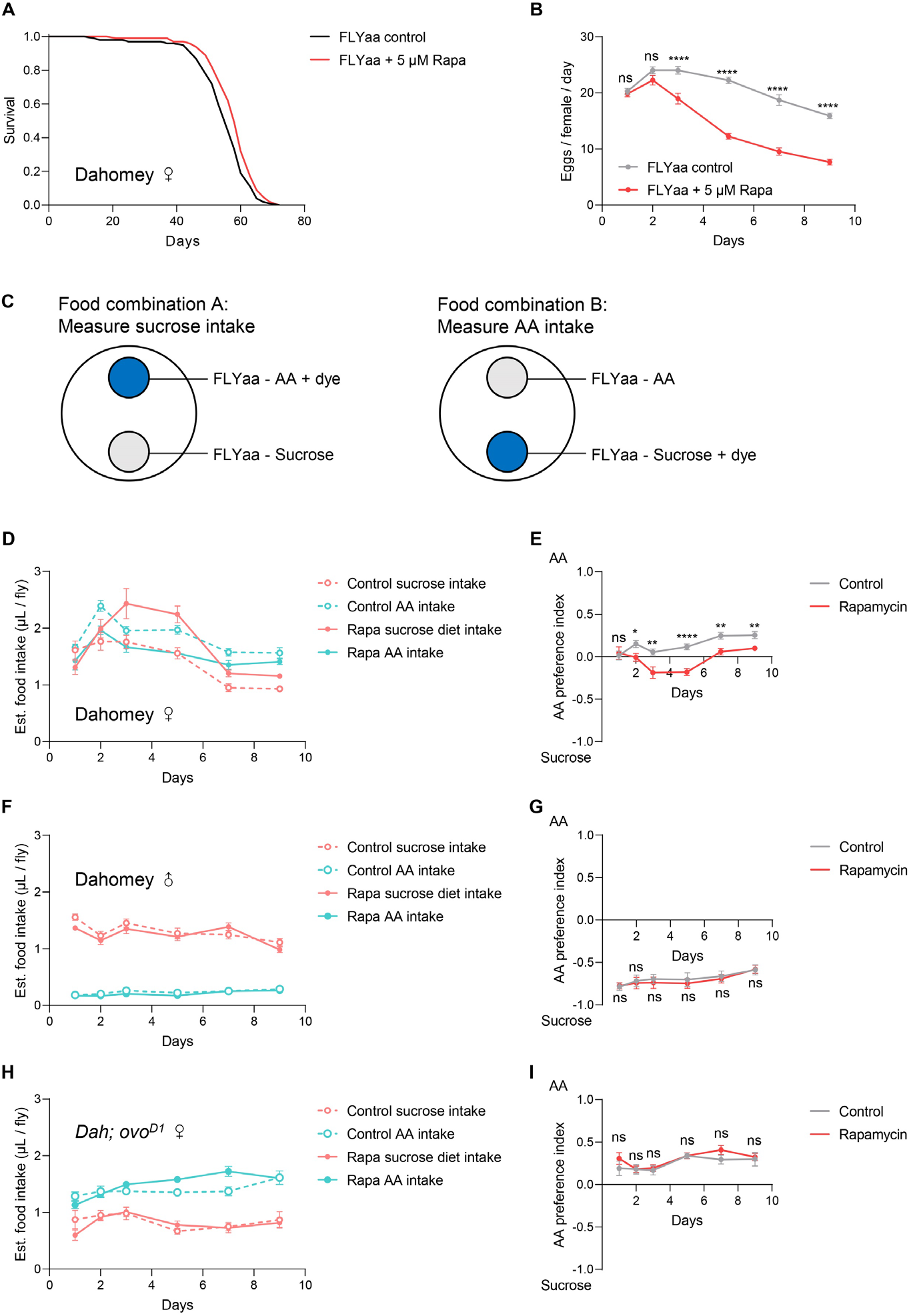
Sex and reproductive dependence of the effects of rapamycin on feeding preference. (A-B) Rapamycin significantly extended the lifespan (A) of wild-type females and decreased their fecundity (B) in FLYaa medium. (C) Schematic diagram of a binary choice EX-Q feeding preference test. In two groups, amino acids-omitted FLYaa medium and sucrose-omitted FLYaa medium were labeled with dye, respectively. (D-E) Rapamycin decreased the amino acid feeding preference of wild type females. (F-I) Rapamycin had no effect on the feeding preferences of males (F-G) and *ovo*^*D1*^ infertile females (H-I).

## 4. Discussion

Previous studies have shown that mTOR signaling pathway is involved in the regulation of feeding preference in *Drosophila*, with a significant sex-specific dimorphism observed. Specifically, neuron-specific activation of S6K enhances female preference for yeast, whereas its inhibition diminishes this preference^24^. Contrastingly, in males, both activation and inhibition of TOR/S6K signaling uniformly increase yeast preference^25^, thereby complicating the mechanistic interpretation of mTOR’s influence on feeding behavior. In this study, we uncover the potential effect of rapamycin on feeding behavior, demonstrating that it modulates female preferences for amino acids and sugars in a manner contingent on reproductive status. However, unlike previous studies, we found that short-term rapamycin treatment did not alter the feeding behavior of males. One possible explanation is that the optimal dose of rapamycin for prolonging lifespan does not inhibit the mTOR signaling to the level that sufficient to alter male feeding behavior. Interestingly, rapamycin also did not alter the preference for amino acids and sugars in *ovo*^*D1*^ sterile females, suggesting that its effect on female feeding behavior may be largely dependent on its inhibition of egg production. However, in aged wild-type females exhibiting reduced fecundity, prolonged rapamycin exposure markedly shifts yeast preference, indicating possible long-term effects on feeding behaviors. These effects might be linked to the permanent modifications of ovarian function and energy metabolism instigated in early life stages. Investigating the effects of long-term rapamycin interventions on feeding preferences in males and sterile females could further elucidate the underlying mechanisms.

Our current explanations of the mechanism by which rapamycin regulates animal feeding preference are largely based on the hypothesis that rapamycin alters macronutrient requirements associated with reproduction. It remains uncertain whether rapamycin affects preferences for non-energy nutrients such as vitamins and minerals, which are pivotal in shaping feeding decisions^21,23,26-28^. Should rapamycin prove to alter preferences for these nutrients, a reassessment of its regulatory logic on feeding behavior would be necessitated.

Taste perception plays a crucial role in the nutrition perception and feeding choices of animals, with preferences for macronutrients including proteins, amino acids, sugars, and other micronutrients reliant on distinct taste neurons^29,30^. Hence, a pertinent question arises as to whether rapamycin-induced feeding behavior changes depends on taste perception, or whether post-ingestion feedback mechanisms, independent of taste, suffice in mediating these effects. Testing the effect of rapamycin on feeding preference in taste-deficient mutant flies will be instrumental in addressing this question.

Overall, our study reveals a previously unrecognized side effect of rapamycin in altering feeding preferences, underscoring potential risks associated with its use as an anti-aging therapeutic.

## Acknowledgments

This work was supported by National Natural Science Foundation of China (32200383) and Natural Science Foundation of Sichuan Province (2022NSFSC1621).

## Author contributions

QW, and GY conceived the experiments; GY, QY and QW performed experiments. All authors analyzed the data. GY, and QW wrote the manuscript. All authors provided final approval of the submitted version.

## Reference

1 Bjedov, I. et al. Mechanisms of life span extension by rapamycin in the fruit fly Drosophila melanogaster. Cell metabolism 11, 35–46 (2010).

2 Li, Z. et al. Aging and age-related diseases: from mechanisms to therapeutic strategies. Biogerontology 22, 165–187 (2021).

3 Kirkland, J. & Tchkonia, T. Senolytic drugs: from discovery to translation. Journal of internal medicine 288, 518–536 (2020).

4 Partridge, L., Fuentealba, M. & Kennedy, B. K. The quest to slow ageing through drug discovery. Nature Reviews Drug Discovery 19, 513–532 (2020).

5 Powers, R. W., Kaeberlein, M., Caldwell, S. D., Kennedy, B. K. & Fields, S. Extension of chronological life span in yeast by decreased TOR pathway signaling. Genes & development 20, 174–184 (2006).

6 Wilkinson, J. E. et al. Rapamycin slows aging in mice. Aging cell 11, 675–682 (2012).

7 Johnson, S. C., Rabinovitch, P. S. & Kaeberlein, M. mTOR is a key modulator of ageing and age-related disease. Nature 493, 338–345 (2013).

8 Selvarani, R., Mohammed, S. & Richardson, A. Effect of rapamycin on aging and age-related diseases—past and future. Geroscience 43, 1135–1158 (2021).

9 Bitto, A. et al. Transient rapamycin treatment can increase lifespan and healthspan in middle-aged mice. elife 5, e16351 (2016).

10 Aiello, G. et al. Transient rapamycin treatment during developmental stage extends lifespan in Mus musculus and Drosophila melanogaster. EMBO reports 23, e55299 (2022).

11 Yang, S.-B. et al. Rapamycin induces glucose intolerance in mice by reducing islet mass, insulin content, and insulin sensitivity. Journal of molecular medicine 90, 575–585 (2012).

12 Lamming, D. W. et al. Rapamycin-induced insulin resistance is mediated by mTORC2 loss and uncoupled from longevity. science 335, 1638–1643 (2012).

13 Kaymakcalan, M. et al. Risk of infections in renal cell carcinoma (RCC) and non-RCC patients treated with mammalian target of rapamycin inhibitors. British journal of cancer 108, 2478–2484 (2013).

14 Harrison, B. R. et al. Wide-ranging genetic variation in sensitivity to rapamycin in Drosophila melanogaster. Aging Cell, e14292 (2024).

15 Piper, M. D. & Partridge, L. Protocols to Study Aging in Drosophila. Methods in molecular biology (Clifton, N.J.) 1478, 291–302, doi:10.1007/978-1-4939-6371-3_18 (2016).

16 Piper, M. D. W. et al. Matching Dietary Amino Acid Balance to the In Silico-Translated Exome Optimizes Growth and Reproduction without Cost to Lifespan. Cell Metab 25, 1206, doi:10.1016/j.cmet.2017.04.020 (2017).

17 Wu, Q. et al. Excreta quantification (EX-Q) for longitudinal measurements of food intake in Drosophila. iScience 23, 100776 (2020).

18 Münch, D., Goldschmidt, D. & Ribeiro, C. The neuronal logic of how internal states control food choice. Nature 607, 747–755, doi:10.1038/s41586-022-04909-5 (2022).

19 Wu, G. et al. Opposing GPCR signaling programs protein intake setpoint in Drosophila. Cell (2024).

20 Bass, T. M. et al. Optimization of dietary restriction protocols in Drosophila. The Journals of Gerontology Series A: Biological Sciences and Medical Sciences 62, 1071–1081 (2007).

21 Wu, Q., Park, S. J., Yang, M. & William, W. J. Vitamin preference in Drosophila. Current Biology 31, R946–R947 (2021).

22 Yu, G., Liu, S., Yang, K. & Wu, Q. Reproductive-dependent effects of B vitamin deficiency on lifespan and physiology. Frontiers in Nutrition 10 (2023).

23 Chen, Y. D. & Dahanukar, A. Recent advances in the genetic basis of taste detection in Drosophila. Cell Mol Life Sci 77, 1087–1101, doi:10.1007/s00018-019-03320-0 (2020).

24 Vargas, M. A., Luo, N., Yamaguchi, A. & Kapahi, P. A role for S6 kinase and serotonin in postmating dietary switch and balance of nutrients in D. melanogaster. Current Biology 20, 1006–1011 (2010).

25 Ribeiro, C. & Dickson, B. J. Sex peptide receptor and neuronal TOR/S6K signaling modulate nutrient balancing in Drosophila. Current Biology 20, 1000–1005 (2010).

26 Shrestha, B., Aryal, B. & Lee, Y. The taste of vitamin C in Drosophila. EMBO reports 24, e56319 (2023).

27 Lee, Y., Poudel, S., Kim, Y., Thakur, D. & Montell, C. Calcium taste avoidance in Drosophila. Neuron 97, 67-74. e64 (2018).

28 Xiao, S., Baik, L. S., Shang, X. & Carlson, J. R. Meeting a threat of the Anthropocene: taste avoidance of metal ions by Drosophila. Proceedings of the National Academy of Sciences 119, e2204238119 (2022).

29 Freeman, E. G. & Dahanukar, A. Molecular neurobiology of Drosophila taste. Current opinion in neurobiology 34, 140–148 (2015).

30 Chen, Y.-C.D. & Dahanukar, A. Recent advances in the genetic basis of taste detection in Drosophila. Cellular and Molecular Life Sciences, 1–15 (2020).

